# Identification and Characterization of Novel Metagenome Assembled Uncultivated Virus Genomes from Human Gut

**DOI:** 10.1101/2024.05.07.592957

**Authors:** Kanchan Bhardwaj, Niharika, Anjali Garg, Aakriti Jain, Manish Kumar, Manish Datt, Vijay Singh, Sudhanshu Vrati

## Abstract

Metagenomics has revealed an unprecedented viral diversity in human gut although, most of the sequence data remains to be characterized. In this study, we mined a collection of 1090 metagenome assembled “high quality” genomes of human gut viruses. Sequence analysis has revealed eight new species from seven genera of the class, *Caudoviricetes* and nineteen new species from fourteen genera of the ssDNA virus family, *Microviridae.* In addition, four “high quality” genomes were identified, which do not show similarity to sequences present in any of the four major viral databases, NCBI viral RefSeq, IMG-VR, Gut Phage Database (GPD) and Gut Virome Database (GVD). Further, annotation of the “high-quality” genomes and KEGG pathway analysis has identified *antB*, *dnaB*, *DNMT1*, *DUT*, *xlyAB*, *xtmB* and *xtmA* as the most widespread viral and Auxiliary Metabolic Genes (AMGs). Genes for virulence, host-takeover, drug resistance, tRNA, tmRNA and CRISPR elements were also found. Bacterial hosts are predicted for around 40% of the analyzed genomes. Overall, we report identification of new viral genomes and genome analyses of human gut viruses, which will be useful for biological characterization to establish their significance in physiology.

**IMPORTANCE:** Multiple studies have found that dysbiosis of gut virome is associated with conditions such as metabolic syndromes, autoimmune disorders and infectious diseases. In the interest of its therapeutic and diagnostic potential, intestinal virome warrants detailed investigation. However, limited ability to culture gut viruses becomes one of the challenges for their biological characterization and fully understanding their role in physiology. Sequence analysis and host prediction methods provide opportunities to understand gut viruses, their functional potential and devise ways for further characterization.

## INTRODUCTION

Application of metagenomics has revealed an unprecedented viral abundance and diversity that is present in nature^1–6^. It has also changed some of the practices in virology such as the way of how viruses have been classified historically^7^. Viruses identified through metagenomics are now being included into the ICTV taxonomy of viruses. For instance, *Genomoviridae* family of ssDNA viruses is populated with sequences obtained by metagenomics^8^. Analysis of invertebrate RNA viral sequences has discovered more than a thousand viruses, provided significant insights into fundamental viral processes and contributed to filling up gaps in the RNA virus phylogeny^9^. Similarly, several other viruses have been discovered through metagenomics, in various ecosystems^10–19^. Since the discovery of crAss-like phages in 2014, as highly abundant dsDNA phages in the human gut metagenomes, there is significant progress in their characterization including the identification of their host^16,20^. Further, analysis of the assembled genomes from metagenomics data, has provided significant insights into structural and mechanistic aspects of viral genomes^21–25^. Impact of virus-encoded metabolic genes on biogeochemical processes and biotechnological applications is revealed through such analyses^26–32^. Correlation of host-associated viruses with host physiology and diseases has also been noted^33–34^. Furthermore, since viruses are old and have dynamic genomes, their origins and evolution could be understood through comparative analyses of large number of sequences obtained through metagenomics. Thus, discovery and analysis of novel virus sequences are of interest for multiple reasons.

In this study, we analyzed the assembled viral genomes that were obtained through metagenome sequencing of total microbial DNA as well as ssDNA and dsDNA extracted from the free virion particles, present in the human fecal samples^35^. We report identification of new viral genomes, new viral species of the class *Caudoviricetes* and family *Microviridae*, the most widespread genes and Auxiliary Metabolic Genes associated with human gut viruses, important structural features of the recovered genomes and their predicted hosts.

## RESULTS

### 1. Identification of novel metagenome assembled uncultivated virus genomes (MAUViG)

We analyzed a collection of 61,099 sequences that were assembled in a previous metagenomics study of human gut viruses (Bhardwaj *et al.*, 2022)^35^. They were grouped into two categories: “high-quality” Metagenome Assembled Uncultivated Viral Genomes (MAUViG) (>90% complete) and “Genome-fragments” (<90% complete or of unknown completeness) with the tool CheckV^36^. CheckV assigns completeness based on comparison of query sequences with a large database of diverse viral genomes (24,834 viral genomes) and examination of the genome ends for terminal repeat sequences, prophage integration sites and host sequences. Based on AAI- or HMM-based approaches, it also assigns confidence level for complete genomes^36^. Among the 1090 “high-quality” sequences, around 80% (874 sequences) were assigned as complete genomes and the rest (216 sequences) were >90% complete. Length of these genomes ranged from 2,285 nt to 720,556 nt and from 3,405 nt to 534,039 nt for the complete and >90% complete genomes, respectively (Figure 1A). For their identification, we performed a search using the NCBI BLASTn program and found that 788 of the sequences were not represented in the NCBI viral RefSeq database. Although, most of them showed presence in the GVD and IMG-VR databases, which was not surprising because these two databases contain data from large number of metagenome studies (Figure 1B). Strikingly, four of the genomes were not found in any of the four major viral databases (Table 1). Notably, all four of these sequences were identified with the virus detection tools, VirFinder and DeepVirFinder, which are trained to identify prokaryotic viruses by k-mer based (deep)machine learning methods. Being homology-independent methods, they are known to have higher potential to identify novel sequences^37^. Among the 788 genomes that remained unaligned to the NCBI viral RefSeq database, 632 were complete genomes and 156 were >90% complete (Figure 1C). The length of the smallest genome was 3, 919 nt (k141_248128_Bac_F21) and that of the largest was 59, 4315 nt (k141_271162_Bac_F29), both of which were identified as complete genomes (Figure 1C). The largest 59,4315 nt (k141_271162_Bac_F29) genome was identified as a provirus with proviral length of 53,088 nt, based on the analysis with the tool, CheckV^36^.

**Figure 1.**
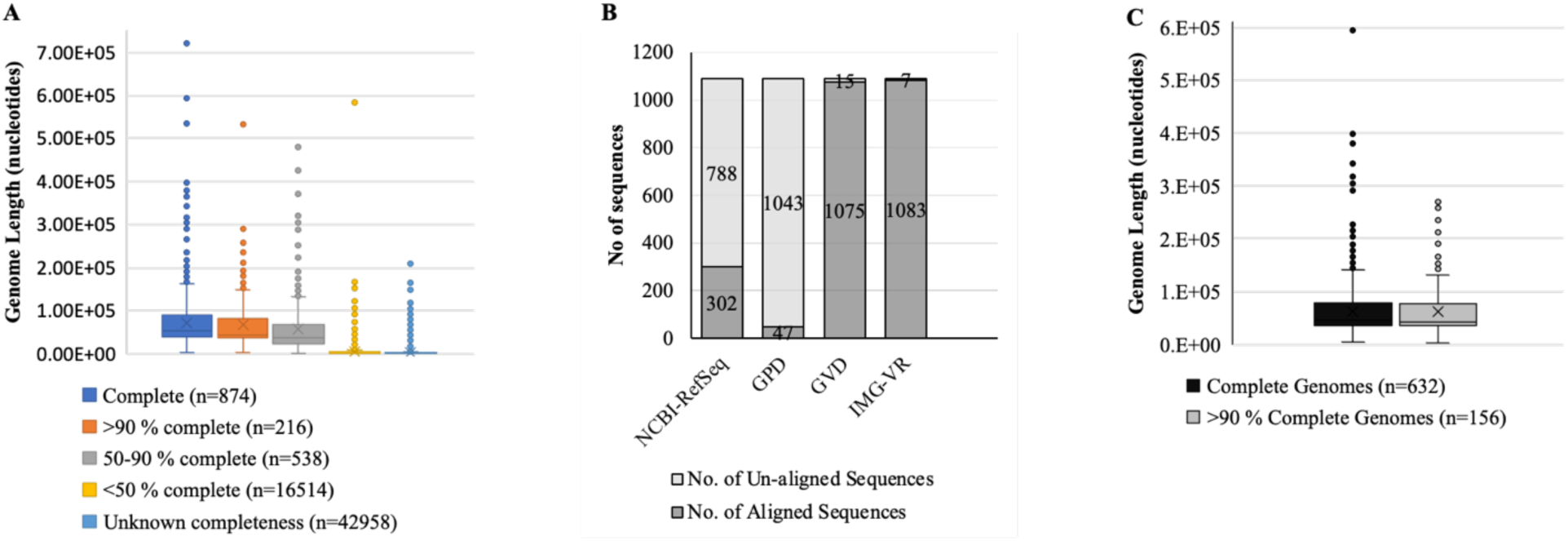
Identification of “high-quality” Metagenome Assembled Uncultivated Viral Genomes (MAUViG). **A.** A total of 61,099 sequences, assembled from metagenome data, were analyzed by CheckV. Their distribution based on genome completeness and length are shown. **B.** Proportion of the “high-quality” MAUViGs (n=1090) that aligned to the four viral databases, NCBI RefSeq, Gut Phageome Database (GPD), Gut Virus Database (GVD) and IMG-VR are shown in dark gray and the number of the unaligned is in lighter shade of gray. **C.** Size distribution of the 788 “high-quality” MAUViGs that remained unaligned to the NCBI RefSeq database.

**Table 1.** Summary of the four novel MAUViGs.

| S. No. | Genome ID | Sample Recovery Methods | Virus Detection Tools | Length (Gene density) | MIUVIG quality | Genome completeness (CheckV) | Completeness method | Lysogeny genes detected |
| --- | --- | --- | --- | --- | --- | --- | --- | --- |
| 1. | K119_1256_SwiBio_F_0_11 | Virus enrichment | VirFinder<br>DeepVirFinder | 13, 358 nt (88.5) | High-quality | 97.4 % | AAI-based (medium-confidence) | Yes |
| 2. | K141_24320_SwiBio_F_0_13 | Virus enrichment | VirFinder<br>DeepVirFinder | 3, 405 nt (96.6) | High-quality | 95.05 % | AAI-based (medium-confidence) | No |
| 3. | K141_37085_Bac_F_0_13 | Without enrichment | VirFinder<br>DeepVirFinder | 5, 674 nt (95.4) | High-quality | 100 % | AAI-based (medium-confidence) | No |
| 4. | K141_248128_Bac_F_0_21 | Without enrichment | VirFinder<br>DeepVirFinder | 3, 919 nt (90.6) | High-quality | 100 % | DTR (medium-confidence) | No |

### 2. Identification of Viral Taxa

For taxonomic identification of the 788 “high-quality” genomes, we annotated them using three of the reported tools, Kaiju, vCONTACT2, and Inphared^38–40^. Kaiju, which utilizes protein alignment method and the RefSeq database, annotated the input 788 genomes into four realms of viruses, including Duplodnaviria (n=674); Monodnaviria (n=33); Varidnaviria (n=16); Riboviria (n=1). Sixty four genomes remained as unclassified viruses and one was identified as cellular DNA. Duplodnaviria included class *Caudoviricetes* (n=624), families *Herelleviridae* (n=25), *Autographviridae* (n=18), *Demerecviridae* (n=2), *Lilyvirus* (n=4) and unclassified genomes (n=64). Monodnaviria included a single family *Microviridae* (n=33) and the realm, Varidnaviria included four families, *Phycodnaviridae* (n=4), *Mimiviridae* (n=9), *Ascoviridae* (n=2), and *Sphaerolipoviridae* (n=1) (Figure 2A). Identification of cellular DNA is likely due to contamination with host DNA and of Riboviria realm due to mis-assignment because in this study, data were collected by sequencing of DNA. For further validation, we also used two additional tools, vCONTACT2 and Inphared. Each tool is based on different algorithms. vCONTACT2 identified the input genomes as the members of viral class *Caudoviricetes* (n=699), families *Microviridae* (n=51, *Phycodnaviridae* (n=3), and other orders (n=35) (Figure 2B). Three hundred eighty three members of the class *Caudoviricetes* and 33 of *Microviridae* family were identified by both Kaiju and vCONTACT2. The third method, Inphared, identified 42 sequences of class *Caudoviricetes* and *Microviridae* family (Figure 2C). Upon comparison, we found that 30 genomes were assigned the same taxonomy by all three methods. Nine of these belong to the class *Caudoviricetes* and twenty one to *Microviridae* family. All of the identified *Microviridae* genomes were complete genomes whereas, seven of the *Caudoviricetes* class members were complete genomes, one was 96.6% complete and another one was 90.6% complete. The genome sizes ranged between 34,241 and 72,973 nt for the *Caudoviricetes* class members and 4,974 and 8,901 nt for the *Microviridae* family members (Figure 2D). The smallest complete genome among our collection of sequences (k141_248128_Bac_F21) could not be assigned a taxonomic identity whereas, the largest one (k141_271162_Bac_F29) was identified as a dsDNA virus of the class *Caudoviricetes*. It is to be noted that the nomenclature of the morphology-based families in the former virus taxonomy system including, *Siphoviridae, Myoviridae, Podoviridae* and the order Caudovirales has recently been replaced with the class *Caudoviricetes*^41^.

**Figure 2.**
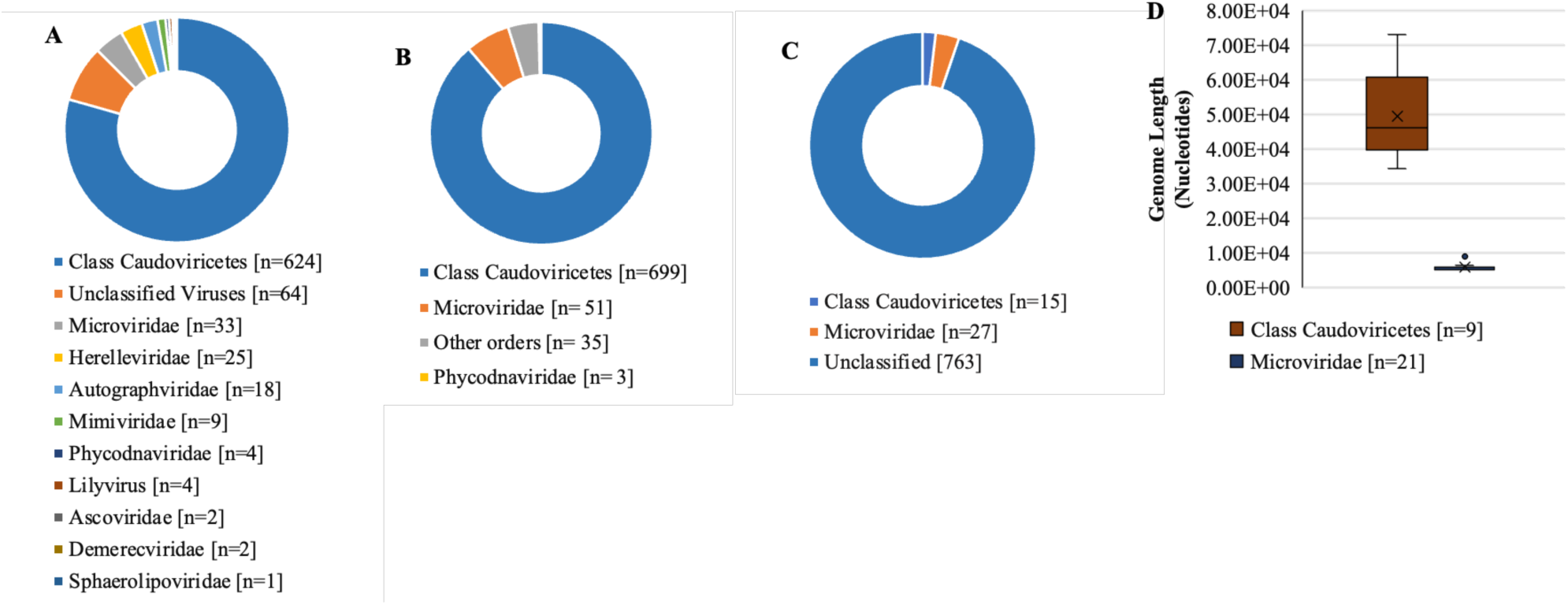
Taxonomic identification of the 788 “high-quality” MAUViGs. **A**. Viral families assigned by the Kaiju method. **B**. Viral families assigned with vCONTACT 2. **C.** Viruses annotated by Inphared. **D.** Size distribution of the genomes that were identified as members of the ds class *Caudoviricetes* and *Microviridae* by all three methods.

### 3. New species of the class *Caudoviricetes*, family *Microviridae* and unidentified taxa

Program, VIRIDIC (Virus Intergenomic Distance Calculator), can be used to determine percent similarity between sequences^42^. As per the genome annotation guidelines of ICTV, if the sequences exhibit >95% similarity (coverage multiplied by identity), they might represent a new strain. Whereas, if the sequence similarity is between 70% and 95%, it might be the 1^st^ isolate of a new species of existing or an undefined genus^43^. VIRIDIC analysis of the thirty genomes that were identified as members of class *Caudoviricetes* and *Microviridae* family by the three tools in our above-mentioned analysis showed that among the nine *Caudoviricetes* phages, the intergenomic similarity of three of them (k141_181068_Bac_F_0_14, k141_49982_SwiBio_F_0_14 and k141_79271_Bac_F_0_21) was between 70% and 95% (Figure 3A). Notably, the completeness of the genome, k141_49982_SwiBio_F_0_14 as predicted by CheckV is 90.6%, while the other two sequences are complete genomes. Manual inspection of the nucleotide sequence alignment showed that k141_49982_SwiBio_F_0_14 was a truncated version of the genome k141_181068_Bac_F_0_14. Further, k141_181068_Bac_F_0_14 and k141_79271_Bac_F_0_21 represent two new species of the same genus in the class *Caudoviricetes*. Whereas, other six genomes likely belong to separate genera in the class *Caudoviricetes* since the intergenomic similarity among them is much lower and ranges from around 0-30% (Figure 3A). Among the genomes of the *Microviridae* family, there were three clusters showing similarity between 70% - 90% and two were 100% similar (Figure 3B). Similarity displayed by the other nine genomes was very low (<50%). Overall, VIRIDIC analysis revealed eight new species of the dsDNA virus class *Caudoviricetes*, which belong to 7 different genera and nineteen new species of the ssDNA virus family, which belong to fourteen separate genera.

**Figure 3.**
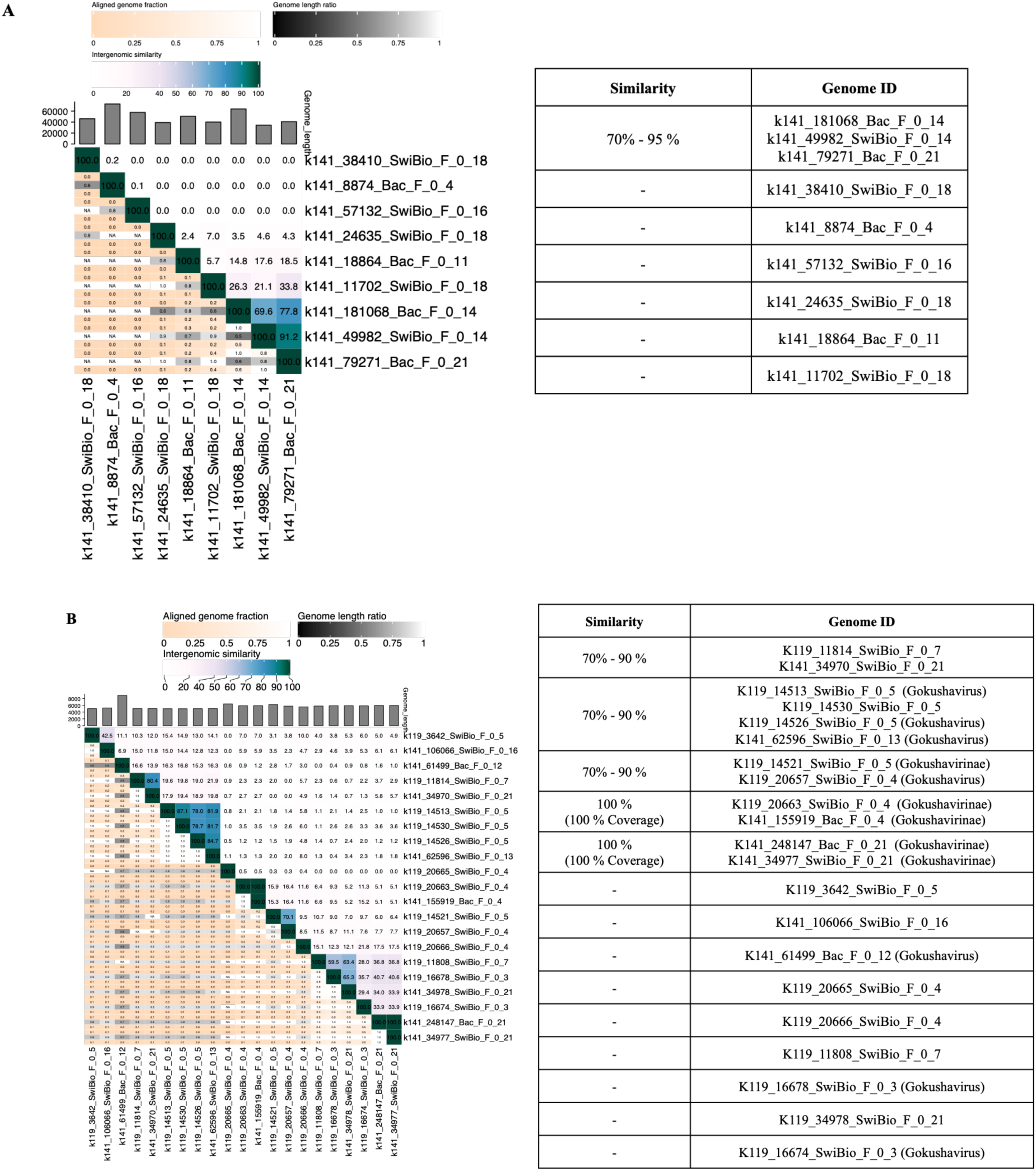
Heatmaps generated by VIRIDIC. The intergenomic similarity values are shown on the right half (upper triangle) and alignment indicators are on the left half (lower triangle). The heatmap is annotated with three different values - aligned genome fraction, genome length ratio, and intergenomic similarity as per the color bars above the plot. **A.** The 9 identified genomes of the class *Caudoviricetes* are listed and the distance among them is indicated in the table. **B.** The 21 identified *Microviridae* genomes and the distance among them is represented and summarized in the table.

In order to understand the relationship of the four “novel” genomes (found absent in all four viral databases) with known phage sequences, we annotated them individually, using the pipeline provided by the Center for Phage Technology at Texas A&M (CPT)^44^. As the first step, CDSs/genes were predicted followed by their annotation using five databases, BLAST SwissProt, BLAST NR (Phages only), InterProscan, KEGG and PFAM. The annotated ORFs were then used for phylogenetic analysis, using NCBI BLASTX search against the non-redundant (nr) database. Five ORFs of the genome, k119_1256_SwiBio_F11, showed a relationship with respective proteins of *Caudoviricetes* and two of them with *Inoviridae* sp. (Figure S1). For the genome, K141_24320_SwiBio_F_0_13, out of the five annotated genes, three were related to the proteins of *Caudoviricetes* and *Inoviridae* sp. (Figure S2). Similarly, the Rep protein of the genome, K141_37085_Bac_F_0_13, showed relationship with *Caudoviricetes* and *Inoviridae* sp. (Figure S3). Whereas, genome, K141_248128_Bac_F_0_21, was related to *Caudoviricetes* sp., CrAss-like virus, *Microviridae* and Bacteriophage sp. (Figure S4). The results are summarized in the Table 2. The other structural features of these genomes, as annotated by the CPT workflow, are summarized in Table 3.

**Figure 4.**
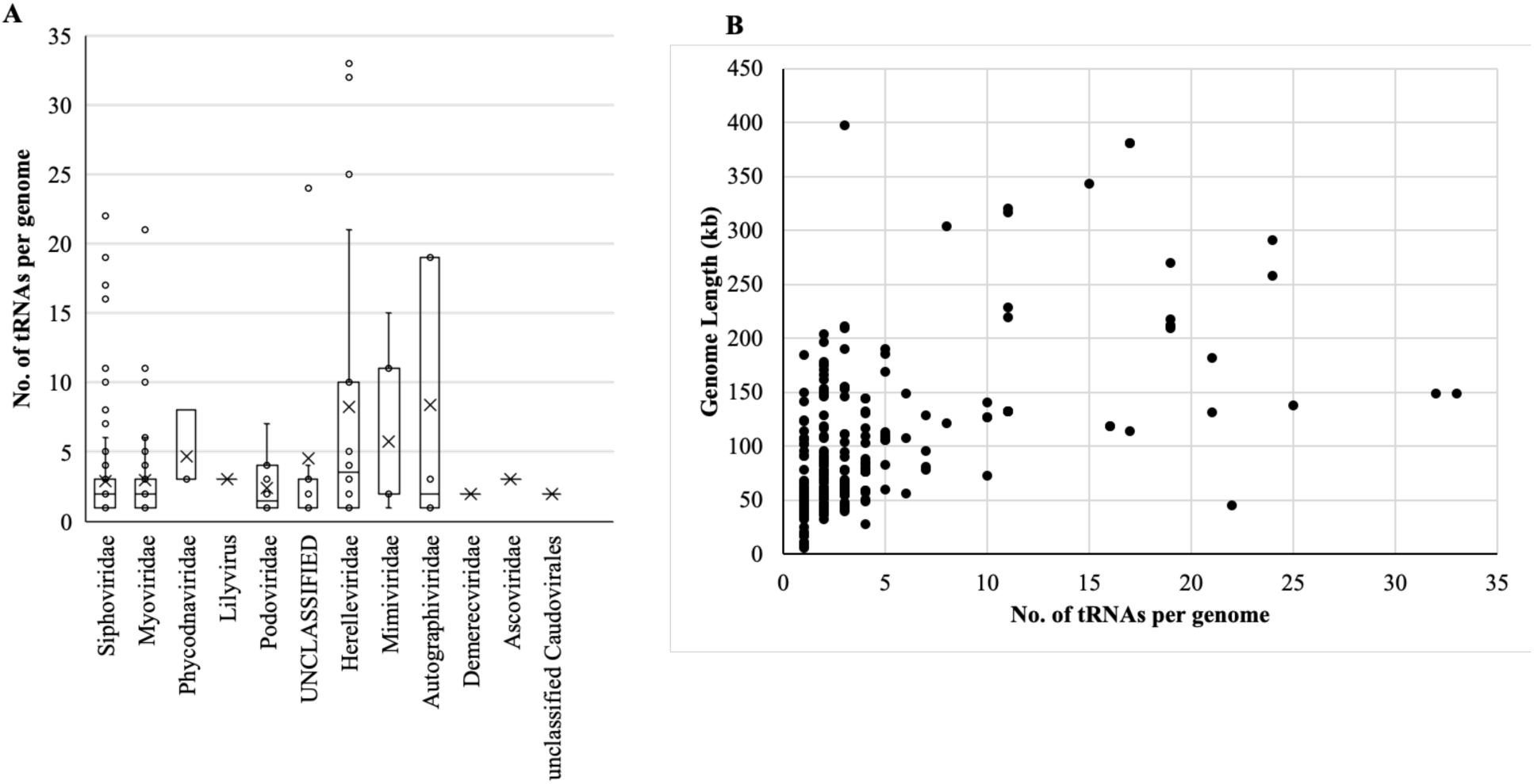
tRNA genes in the 788 “High-quality genomes”. **[A]** Number of tRNA genes found in the identified viral families. **[B]** Correlation between genome length and number of tRNAs.

**Table 2.** Summary of the phylogenetic analysis of the four MAGs.

| Genome ID | ORF/Gene for phylogenetic analysis | Annotation by BLAST SwissProt/BLAST NR (Phages only)/InterProscan/KEGG/PFAM | Identified species through NCBI-BLASTX search |
| --- | --- | --- | --- |
| **K119_1256_SwiBio_F_0_11**<br>(Genome Length= 13, 358 bp )<br>(No. of predicted ORFs = 24) | ORF 4 | Helicase/ATPase/DNA dependent DNA polymerase | *Caudoviricetes* sp. |
|  | ORF 10 | CI repressor/putative antitoxin | *Caudoviricetes* sp. |
|  | ORF 20 | Recombinase/Resolvase/Integrase | *Caudoviricetes* sp., *Inoviridae* sp. |
|  | ORF 21 | Hypothetical protein/PNG 27/SHD of GIT/Nus B family protein | *Caudoviricetes* sp., *Inoviridae* sp. |
|  | ORF 24 | Recombinase/Resolvase/Hypothetical protein | *Caudoviricetes* sp. |
| **K141_24320_SwiBio_F_0_13**<br>(Genome Length= 3, 405 bp )<br>(No. of predicted genes=5) | Gene 2 | Hypothetical protein | Bacteriophage sp. |
|  | Gene 4 | Rep protein | *Caudoviricetes* sp., *Inoviridae* sp. |
|  | Gene 5 | DNA Translocase FTSK | *Caudoviricetes* sp. |
| **K141_37085_Bac_F_0_13**<br>(Genome Length= 5, 674 bp)<br>(No. of predicted ORFs = 9) | ORF 9 | Rep Protein | *Caudoviricetes* sp., *Inoviridae* sp. |
| **K141_248128_Bac_F_0_21**<br>(Genome Length= 3, 919 bp)<br>(No. of predicted genes = 3) | Gene 1 | Plasmid recombination enzyme, Hypothetical protein | *Caudoviricetes* sp., CrAss-like virus, *Microviridae*, Bacteriophage sp. |

**Table 3.** Genome annotation of the four novel MAGs with CPT Workflow.

| Genome ID | tRNA/<br>tmRNA | No. of<br>Terminators<br>(minimum<br>score = 90) | Inverted repeats/<br>Palindromes<br>*Minimum length of palindrome=10<br>*Maximum length of palindrome=100<br>*Maximum gap b/w repeated sequences=100<br>*Number of allowed mismatches = 0 | Tandem Repeats<br>(start-end)/ size/ count<br>*Threshold 10<br>*Min repeat: 10<br>*Max repeat: 10<br>*Mismatch: No | CpG islands<br>*Window size: 10<br>*Calculated % composition of G+C = over 50 %<br>*Calculated Obs/Exp ratio= over 0.6<br>*Conditions should hold for a minimum of 200 bases |
| --- | --- | --- | --- | --- | --- |
| K119_1256_SwiBio_F_0_11<br>(Genome Length= 13, 358 bp ) | Not<br>Found | 08 | 92...101& 163...172;<br>1290...1299& 1300...1309 (PALINDROME);<br>2312...2321& 2382...2391;<br>2321...2332& 2415...2426;<br>2504...2514& 2523...2533;<br>2321...2332& 2349...2360;<br>9487...9497& 9514...9524;<br>11856...11865& 11876...11885 | (1335-1387)/ 10/ 5<br>(1581-1643)/ 11/ 5<br>(2347-2435)/ 22/ 4 | 1433...1775;<br>2627...3594;<br>3852...4221;<br>4662...4959;<br>5126...5407;<br>7334...7670;<br>7867...8097;<br>9072...9306;<br>10288...10715;<br>10799...11539;<br>12057...12425;<br>12772...13301 |
| K141_24320_SwiBio_F_0_13<br>(Genome Length= 3, 405 bp ) | Not<br>Found | 03 | 1826..1835 & 1845..1854;<br>2608..2618 & 2629..2639;<br>2676..2687 & 2688..2699 (PALINDROME) | No tandem repeats<br>found | 855..1063;<br>1400..1756;<br>3046..3279 |
| K141_37085_Bac_F_0_13<br>(Genome Length= 5, 674 bp) | Not<br>Found | 09 | 913...927 & 929...943;<br>1763...1778 & 1784...1799;<br>2770...2779 & 2833...2842;<br>5593..5602 & 5617...5626 | No tandem repeats<br>found | 325...537;<br>2749...2967 |
| K141_248128_Bac_F_0_21<br>(Genome Length= 3, 919 bp) | Not<br>Found | 01 | 1374..1384 & 1403..1413;<br>1708..1718 & 1723..1783;<br>3405..3415 & 3457..3467;<br>3426..3439 & 3506..3519;<br>3702..3713 & 3751..3762 | (762-824)/ 21/ 3 | No CpG islands found |

### 4. Genome annotation

To identify genome features of the 788 sequences, we selected the Pharokka pipeline to detect the coding sequences (CDS), clustered regularly interspaced short palindromic repeats (CRISPRs), transfer RNAs (tRNAs), transfer-messenger RNAs (tmRNAs) antibiotic resistance and virulence genes^45–51^. We detected tRNA genes in 332 out of the 788 sequences. There were up to 33 tRNA genes in a single genome of the *Herelleviridae* family and no tRNA genes were detected in the phages of the *Microviridae* and *Sphaerolipoviridae* families (Figure 4A). No correlation was observed between the number of tRNA genes and the genome length (Figure 4B). In addition, among the 788 sequences, CRISPR repeat sequences were found in two, tmRNA in seven, and virulence genes in two of them (Table 4).

**Table 4.**
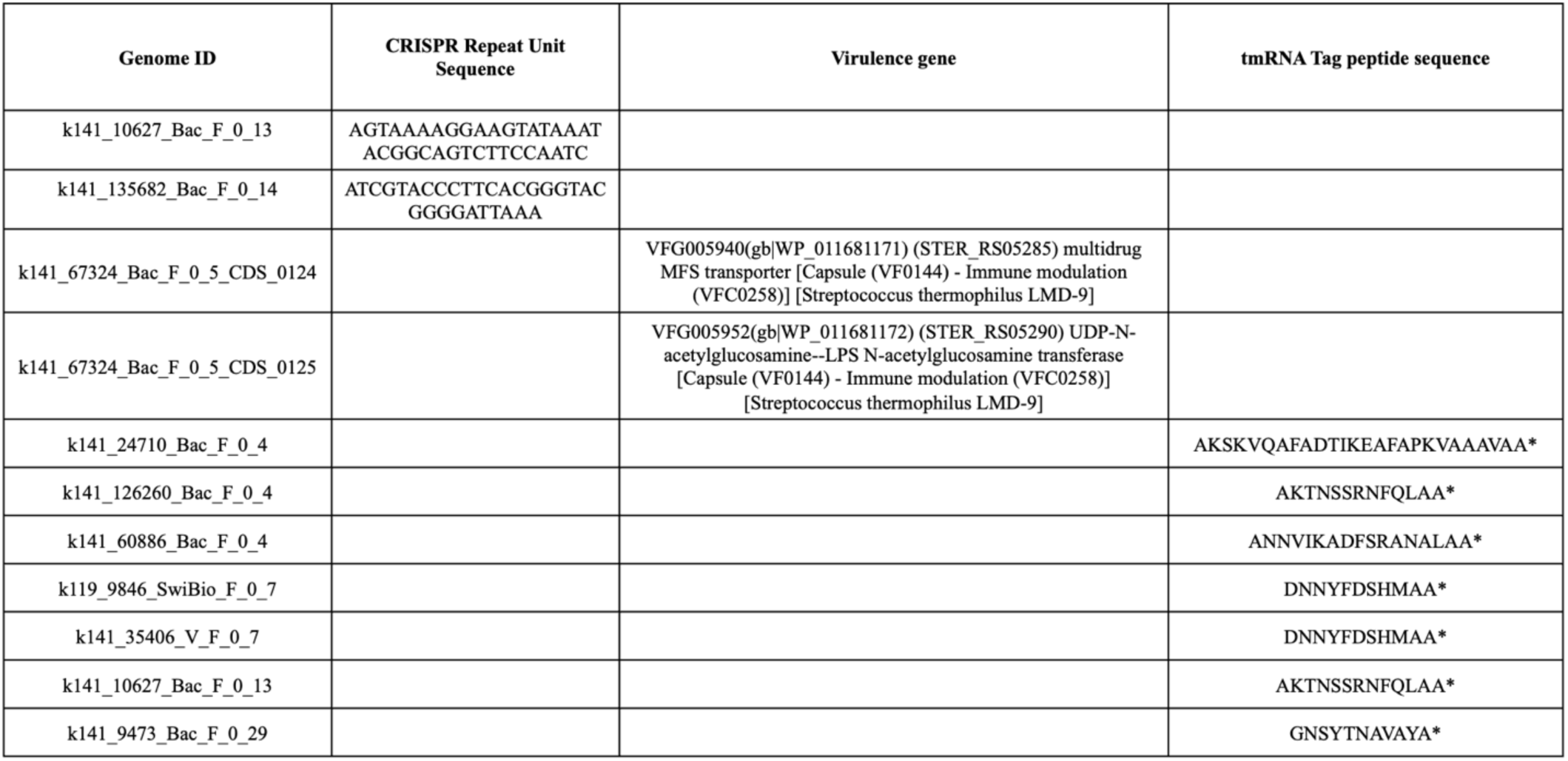
Annotation of the 788 “High-quality genomes”.

### 5. Functional analysis

First, we used the Pharokka pipeline for functional annotation of the genomes because it uses Phanotate to predict CDS and Phanotate is known to predict significantly higher number of CDS than other gene prediction tools as it considers the unique features of phage genomes such as the alternative start codons, high coding density and small gene size^45,51^. Pharokka pipeline uses PHROGs database for annotation of the predicted CDSs. PHROGs database contains 868,340 proteins from 17,473 genomes of phages of bacteria or archaea, which are grouped into 38,880 PHROGs (protein orthologous groups)^52^. All of the predicted CDS in our collection of 788 “High-quality genomes”, upon translation into a protein sequence and assignment to the closest protein in the PHROG database, were grouped into following nine classes: Connector; DNA, RNA and Nucleotide Metabolism; Head and Packaging; Integration and Excision; Lysis; Moron, Auxilliary Metabolic Gene and Host Takeover; Tail; Transcription Regulation and Others (Figure 5A). Notably, more than 80% of the CDS could not be assigned functions. All of the viral families (identified as per the Kaiju classification) encoded the genes for viral functions but only 312 genomes, representing 8 viral families and UNCLASSIFIED viruses contained CDS for one or more of the 52 “Moron, Auxilliary Metabolic Gene and Host Takeover proteins” (Figure 5B). Although most of the genomes had a single “Moron, Auxilliary Metabolic Gene and Host Takeover proteins”, up to 11 such genes were detected in a single genome. Among the eight novel species of the class *Caudoviricetes*, only one (K141_8874_Bac_F_0_4) had a “Moron, Auxilliary Metabolic Gene and Host Takeover proteins” gene, phosphoadenosine phosphosulfate reductase, and one of the four novel genomes (K119_1256_SwiBio_F_0_11) contained a gene encoding HicB-like toxin-antitoxin system. None of the nineteen novel species of *Microviridae* family had any “Moron, Auxilliary Metabolic Gene and Host Takeover proteins”.

**Figure 5.**
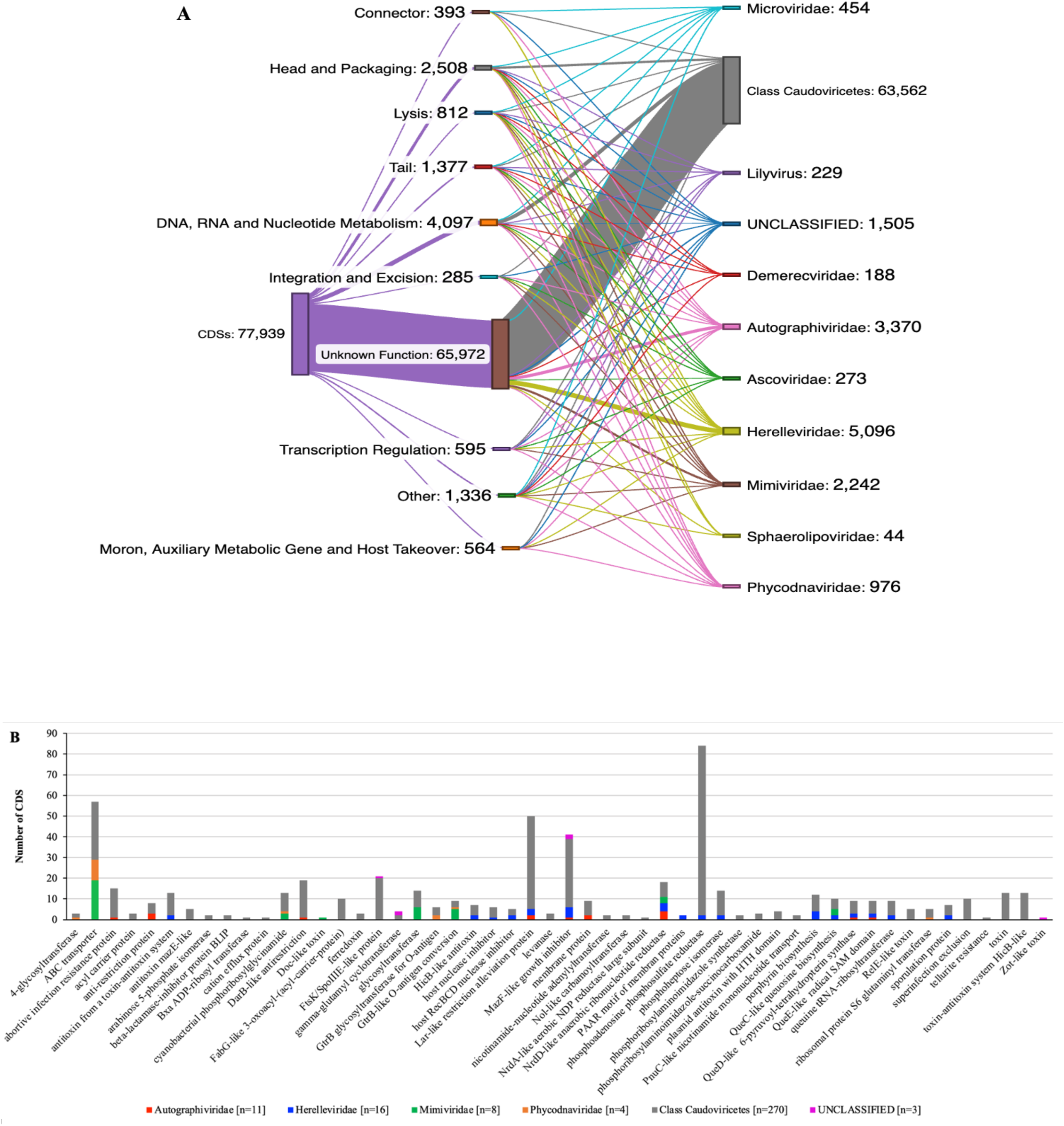
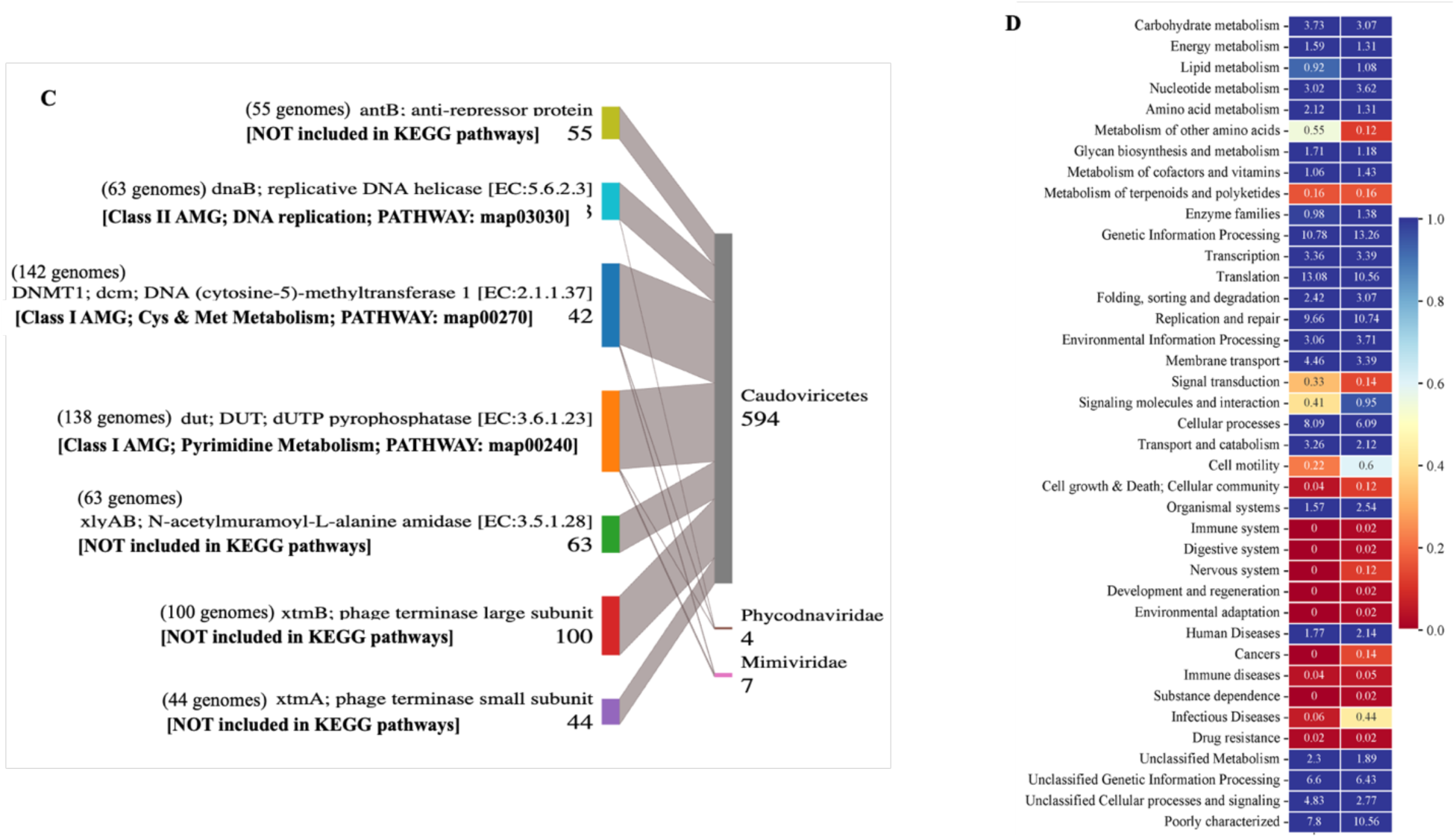
Functional annotation of the 788 “High-quality genomes”. **[A]** Sankeymatic diagram showing proportion of the nine identified functional categories and the CDSs with unknown functions. Number of identified CDS in each category and the number of CDS for which functions could not be identified are given. Viral families (as per Kaiju annotation) corresponding to each functional category and the total number of CDS identified in each family are shown on the right. [**B] Distribution of the CDSs, encoding “Moron, Auxiliary Metabolic Genes and Host Takeover” proteins.** Protein functions are denoted on the horizontal axis and the vertical axis shows their proportion found among 11 members of *Autographviridae*, 16 *Herelleviridae*, 8 *Mimiviridae*, 4 *Phycodnaviridae,* 270 class *Caudoviricetes* and 3 Unclassified families. Each viral taxa are indicated by different colors. [**C] Sankeymatic diagram showing proportion of seven core AMGs and corresponding viral taxa.** *antB*: Transcription related protein [Genetic information processing]; *dnaB*: Replication and repair [Genetic information processing]; *DNMT1*: Amino acid metabolism [Metabolism]; *DUT*: Pyrimidine metabolism [Metabolism]; *xlyAB*: Viral protein family; *xtmB*: Replication, recombination and repair proteins; *xtmA*: Replication, recombination and repair proteins. [**D] KEGG Mapping.** Distribution of genes across KEGG pathways and processes is shown for the temperate (T) and lytic (L) phages. Proportion (%) of genes in each functional category is denoted inside the boxes.

Second, we used KEGG pathway mapping for gaining insight into the functional potential of the genomes. Some viruses are known to carry host-derived genes, called Auxiliary Metabolic Genes (AMGs), which have been divided into class I and class II^53^. Class I AMGs encode proteins involved in metabolic functions and are included in the KEGG metabolic pathways. Class II AMGs encode proteins with peripheral functions and are not included in the KEGG metabolic pathways. Through analysis of viral genomes obtained from multiple environments, Kieft *et. al.,* (2020) have identified 14 globally conserved AMGs^54^. Further, Luo *et. al.,* (2022) observed a correlation between viral lifestyle and AMG profiles^55^. We performed KO annotation and KEGG mapping of the 788 “High-quality genomes” with the GhostKOALA tool, to examine the presence of core AMGs and correlation of AMG profiles with viral lifestyle^56^. Our 788 “High-quality genomes” were derived from 12 “healthy” individuals. Therefore, in order to find potential core AMGs, the 12 samples were annotated separately with KEGG databases. This analysis identified presence of seven genes-*antB, dnaB, DNMT1(dcm), dut, xlyAB, xtmB, xtmA*, in every individual (Figure 5C). Two of these core genes are Class I and one is Class II AMG. The other four, antB, xlyAB, xtmA and xtmB, encode proteins required for fundamental viral processes and are not included in KEGG pathways, metabolic or others. The identified Class I AMGs, *DNMT1* (*dcm*) and *dut*, code for enzymes involved in metabolism of amino acids and nucleotides, respectively. The Class II AMG, *dnaB*, encodes replicative DNA helicase enzyme involved in the process of DNA replication, not in a metabolic pathway. All seven core genes were found in the viruses of the class *Caudoviricetes* (Figure 5C). In addition, *DNMT1* and *dut* genes were also found in two families of the nucleocytoplasmic large DNA viruses, *Phycodnaviridae* and *Mimiviridae* (Figure 5C). The AMG, *dnaB* was found in members of the class *Caudoviricetes* and the family *Phycodnaviridae* (Figure 5C).

Next, we sorted the genomes into temperate and lytic viruses. Genomes which contained lysogeny related genes, integrase, excisionase, transposase, recombinase, parA/B and CI/Cro repressor were considered temperate and the rest as lytic. Analysis of their genes showed that genetic diversity was higher in lytic viruses and that the class II AMGs are among the dominant genes in both, temperate phages and lytic phages. In the temperate phages, maximum proportion of genes belong to the processes of translation (13.1%), genetic information processing (10.8%), replication and repair (9.7%), cellular processes and signaling (8.1%), poorly characterized (7.8%), unclassified genetic information processing (7.3%), unclassified cellular processes and signaling (5.1%) and membrane transport (5.5%) followed by the class I AMGs for the process of carbohydrate metabolism (3.7%). Although, the gene distribution profiles across the two lifestyles varies to some extent (Figure 5D). For instance, in the lytic phages, maximum proportion of genes belong to the processes of genetic information processing (13.3%), replication and repair (10.8%), translation (10.6%), poorly characterized (10.6), unclassified genetic information processing (6.4%), cellular processes and signaling (6.1%) and environment information processing (3.7%) followed by the class I AMGs for the process of nucleotide metabolism (3.6%). Notably, this analysis also detected two genes for antibiotic resistance vanW; vancomycin resistance protein tetM; tetO; ribosomal protection tetracyclin resistance protein.

### 6. Host Prediction

Identification of host is an important requirement to be able to culture the novel viruses and their characterization. Since majority of the genomes among our collection were identified as the viruses that infect prokaryotes, we used the tool, Prokaryotic Virus Host Predictor, (PHP) to perform viral host prediction^57^. In addition, hosts were also identified by the Inphared program^40^. Among our 788 “high-quality genomes”, PHP predicted the hosts for 323, which included the dsDNA phages and the ssDNA viruses (Figure 6A). The most frequently identified hosts were of the *Ruminococcaceae* family followed by *Lachnospiraceae, Bifidobacteriaceae, Prevotellaceae, Clostridiaceae*. *Oscillospiraceae, Bacteroidaceae, Atopobiaceae, Selenomonadaceae, Veillonellaceae, Eubacteriaceae, Sporomusaceae, Bacillaceae, Lactobacillaceae, Paenibacillaceae and Flavobacteriaceae* (Figure 6A). Among the novel species of class *Caudoviricetes* and *Microviridae* family identified in this analysis, Bifidobacterium was identified as the host for five of the *Caudoviricetes* species by both, PHP and InPHARED. *Phascolarctobacterium succinatutens* was identified as the host for only one of the *Microviridae* species by PHP. InPHARED identified hosts for all the *Caudoviricetes* species but none of the *Microviridae* species (Figure 6B). For three of the four “novel” genomes, the predicted host species were Butyrate-producing bacterium SS3/4(245014) for k119_1256_SwiBio_F11, *Candidatus Peregrinibacteria* bacterium GW2011_GWA2_47_7(1619058) for k141_24320_SwiBio_F13, and *Lactobacillus murinus(1622)* for k141_37085_Bac_F13 (Figure 6B). Further, hosts were predicted for all members of *Phycodnaviridae* and *Mimiviridae* families. Clostridiales bacterium UBA4706(1950889) was predicted the host species for the single viral member of the family *Sphaerolipoviridae* among our sequences. The hosts for our viruses with the smallest (k141_248128_Bac_F21; 3,919 nt) and largest genomes (k141_271162_Bac_F29; 59,4315 nt) could not be identified.

**Figure 6.**
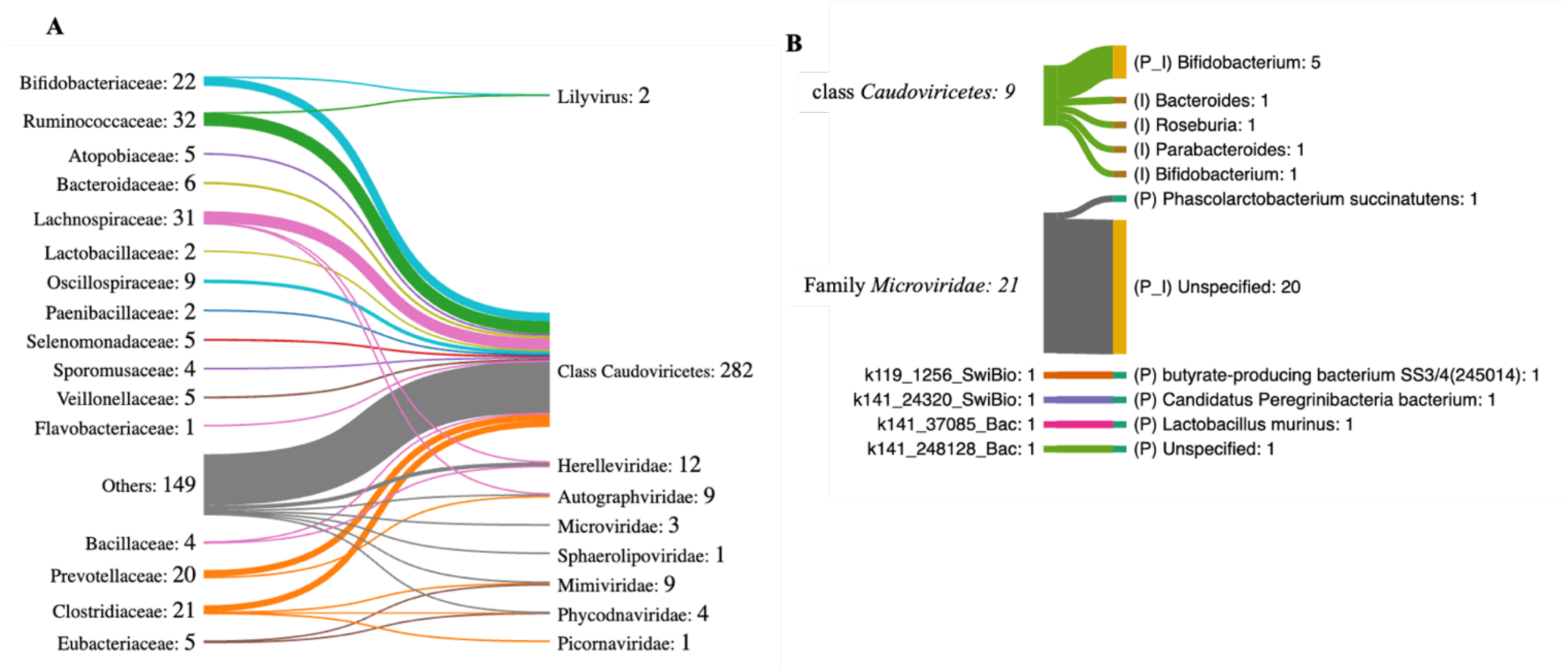
Host Prediction. **[A]** Predicted hosts with PHP. Families of the host bacteria are given on the left and the number of phages for which they are assigned as the host is given next to it. Phage families, identified by Kaiju, are given on the right. The number of phages for which the host could be predicted is mentioned next to the phage family names: Class *Caudoviricetes* (n=282/601); Lilyvirus (2/4); family *Herelleviridae* (n= 12/25); family *Microviridae* (n= 3/33); family *Sphaerolipoviridae* (n= 1/1); family *Autographviridae* (n=9/ 18); family *Phycodnaviridae* (n=4/4) and family *Mimiviridae* (n= 9/9). Hosts could not be identified for the following: Unclassified viruses (n= 0/67); Unclassified members of class *Caudoviricetes* (n=23); family *Ascoviridae* (n=2) and family *Demerecviridae* (n=2). **[B].** Predicted hosts for the newly identified species of the class *Caudoviricetes*, (n=9) and family *Microviridae* (n=21) and the four novel viral genomes. (P) represents the assignment by PHP; (I) by INPHARED and (P&I) by both PHP and InPAHRED.

## METHODS

### Estimating the genome quality and taxonomic annotation of the viral sequences

The tool CheckV was used to estimate the quality of the Uncultivated Viral Genomes (UViGs)^36^. The three modules of CheckV are 1. identify and remove host contamination on proviruses *i.e.* the non-viral genes on the edges of contigs; 2. estimate completeness for genome fragments, based on the expected genome length through either AAI or HMM-based approach; and 3. predict closed genomes based on terminal repeats (ITR, DTR) and provirus integration site. CheckV then classifies sequences into five categories: 1. Complete (DTR, ITR, provirus); 2. High-quality (>90% completeness); 3. Medium quality (50-90% completeness); 4. Low-quality (0-50% completeness); 5. Undetermined quality (completeness estimates not available).

### Genome Annotation

Pharokka (v1.3.2) pipeline was used to detect the coding sequences (CDS) with Phanotate v1.5.1; clustered regularly interspaced short palindromic repeats (CRISPRs) with MinCED (v0.4.2); transfer RNAs (tRNAs) with tRNAscan-SE (v2.0.11); transfer-messenger RNAs (tmRNAs) with Aragorn (v1.2); antibiotic resistance with Comprehensive Antibiotic Resistance Database (CARD) and virulence genes with Virulence Factor Database (VFDB)^45–51^. For the functional annotation of the CDSs predicted by Phanotate (v1.5.1), PHROGs (protein orthologous group) database was used with mmseqs2 tool at a default e-value of 10^-5^. Pharokka was run in the meta mode, which is suitable for metavirome samples. INPHARED was used to identify sequences based on their similarity to INPHARED database of known phages. In addition, CPT Galaxy-Apollo suite was applied for additional annotation of the four novel genomes. Structural annotation was performed by PAP structural workflow v2021.02 of Galaxy-Apollo suite. Additional structural annotations were done using the plugins “palindrome”, “newcpgreport” and “equicktandem”. Functional annotation was performed by PAP Functional workflow v2022.01 of Galaxy-Apollo suite. GhostKOALA was used for KO annotations and KEGG pathway analysis.

### *In silico* host prediction

PHP is a gaussian model based tool and largely falls under the alignment-free methods of host prediction. PHP predict hosts for viral genomes/short contigs based on k-mer signals. It identifies hosts based on consensus of top 30 predictions and has been shown to have an accuracy of 34%, 45%, 60%, 78%, 80% and 98% at genus, family, order, class, phylum and domain levels, respectively^57^. It’s database contains 60,105 prokaryotic genomes as model hosts.

## DISCUSSION

We report identification of eight new species of the dsDNA virus class *Caudoviricetes,* and 19 new species of the ssDNA virus family *Microviridae*. Phages of both these taxa are important gut residents and alteration in their abundance has been associated with diseases^58^. *Microviridae* is one among the five families of ssDNA bacteriophages. Based on the presence or absence of spikes, *Microviridae* family has been divided into two lineages: *Gokushavirinae*, which lacks spikes and Microvirus, which has spikes^59^. The genome size range (4883-6289 nt) and per cent GC content (39.277% - 50.346%) of our sequences is similar to what has been reported for other members of this family of phages^60^. Further, as reported earlier for the *Microviridae* family members, in our analysis also, we did not find Auxilliary Metabolic Genes (AMGs) in any of the *Microviridae* family members identified here. The three most commonly found hosts for *Microviridae* family of bacteriophages belong to the phyla, Proteobacteria, Firmicutes and Bacteroidetes. For a few members, the assigned hosts belong to the phyla: Chlamydiae, Chlorobi, Cyanobacteria/Melainabacteriagroup, Fusobacteria, Margulisbacteria, Melainabacteria, Methanomicrobia, Nitrospirae, Planctomycetes, Spirochaetes, Synergistetes, Tenericutes and Verrucomicrobia^61^. In this analysis, the predicted host for one of the *Microviridae* member is *Phascolarctobacterium succinatutens* sp. from the family *Acidaminococcaceae,* phylum, Firmicutes. Hosts could not be predicted for the other 13 members.

Phages of the class *Caudoviricetes* represent one of the most abundant groups of viruses found in human gut^62,63^. These phages possess dsDNA linear genomes with ends containing Direct Terminal Repeats (DTRs), Inverted Terminal Repeats (ITRs), or host genome fragments^64–66^. Other features of these phages are that they encode distinctive capsid protein and terminase enzyme^67,68^. Lifestyle of these phages could be virulent, temperate or they may exist as cryptic prophages^69,70^. Among the eight novel species of the class *Caudoviricetes* that we identified in this analysis, none possess lysogeny related genes, three of them have DTRs, all encode Terminase large subunit and five were detected with coding sequence for Major Head Protein. Further, none of them have CRISPR sequence or tmRNA genes. Although, one contains two tRNA and another one contains 10 tRNA genes. The four novel genomes could not be assigned a certain viral taxa. However, based on their predicted hosts, we plan to try and culture them for further characterization.

“Mosaic” and “modular” nature are the characteristic features of bacteriophage genomes. In addition to the “hallmark genes”, many phages encode genes that perform non-essential viral functions, referred to as Auxiliary Metabolic Genes (AMGs). Phage encoded AMGs are known to impact bacterial metabolism, adaptability, pathogenicity and colonization^71–74^. In the oceans and soil habitats, some of the virus-encoded AMGs have been characterized and are either shown or implicated to have roles in global processes^75–77^. The 14 globally-conserved AMGs identified by Kieft *et. al.,* (2020) include *ahbD, cobS, cysH, dcm, folE, mec, moeB, phnP, queE, queD, queC, ubiG, ubiE, waaF*^54^. In our analysis of human gut viruses, the most widespread viral AMGs were DNMT1 (*dcm*) encoding DNA (cytosine-5)-methyltransferase I (present in all individuals); DUT (*dut*) encoding dUTP pyrophosphatase (present in all individuals); *dnaB* encoding the replicative DNA helicase (present in all individuals); *cysH*, encoding phosphoadenosine phosphosulfate reductase (present in 83% of the individuals) followed by three genes, NrdD-like anaerobic ribonucleotide reductase, MazF-like growth inhibitor and ABC (ATP-binding cassette) transporter (present in 75% of the individuals) (Figure 5B & 5C). Both, phosphoadenosine phosphosulfate reductase and NrdD-like anaerobic ribonucleotide reductase are enzymes involved in the metabolism of sulfur and nucleotides, respectively. Whereas, MazF-like growth inhibitors represent the toxin molecule of bacterial toxin-antitoxin system^78^. ABC transporters are ubiquitous proteins that catalyze the translocation of their substrates and can be classified as importers, exporters or extruders based on the direction of substrate translocation^79^. Similar to Luo *et al.* (2022), our analysis showed that the gene diversity is higher in the lytic phages as compared to the temperate^55^. Highest proportion of genes in phages with both lifestyles encode functions for genetic information processing, replication and repair, translation, cellular processes and signaling and environment information. Among the class I AMGs, temperate phages showed higher proportion of genes for metabolism of amino acids, carbohydrates, energy and glycan biosynthesis. Whereas, in lytic phages it was for metabolism of nucleotides, lipids and cofactors and vitamins.

Notably, two virulence factors were detected in one of the 1,74,344 nt long complete genomes (k141_67324_Bac_F_0_5). They are multidrug MFS transporter and UDP-N-acetylglucosamine--LPS N-acetylglucosamine transferase. In addition, this genome contained ABC transporter, GtrB glycosyltransferase for O-antigen, GtrB-like O-antigen conversion and ribosomal protein S6 glutaminyl transferase, 237 CDS with unknown function and no genes for tmRNA, tRNA or CRISPR. It has an integrase gene and no lysis genes, indicative of a cryptic prophage (a temperate phage which might have lost the capacity to be lytic). This genome was identified as a member of *Phycodnaviridae* family by Kaiju and *Caudoviricetes* class member by vCONTACT2 (although, no tail protein was predicted). *Eubacterium ventriosum* was identified as the host species for this phage, which is a gram-positive bacterium of the family *Eubacteriaceae* of the order Clostridiales and Phylum Firmicutes. Although, the Major Facilitator Superfamily (MFS) transporter is a listed virulence factor in the Virulence Factor Database, possibly due to their association with the mechanisms of antimicrobial resistance in pathogens, MFS transporter and ATP-binding cassette superfamily transporter (ABC transporter) are important family of ubiquitous membrane transporters involved in active extrusion, which is one of the mechanisms by which cells react when exposed to toxic compounds^80–82^. UDP-N-acetylglucosamine--LPS N-acetylglucosamine transferase catalyzes the first step in the synthesis of Lipid A that makes up the outer LPS layer of the gram-negative bacteria. It is also interesting to detect the genes for GtrB glycosyltransferase for O-antigen and GtrB-like O-antigen conversion because O-antigen is a part of LPS and bacteriophage mediated modification of O-antigen, leading to serotype conversion and alteration of phage receptors, has been reported^83^. Intriguingly, the predicted host for this phage genome is a gram-positive bacterium. Therefore, it is possible that the host prediction is inaccurate. Alternatively, this genome is likely a reservoir for some of the genes which might have been picked up in the process of recombination. RimK (ribosomal protein S6 glutaminyl transferase) is known to modulate the SOS response in *Escherichia coli* and it has been suggested to be involved in the response to oxidative or UVA stress in cyanobacteria^84–85^.

The analysis presented here will facilitate further characterization of some of the viruses and virus-encoded functions. Notable, most of the genomes and genes still remain un-annotated. This again highlights the need for additional tools and techniques for taxonomic and functional annotation of viral sequences. Further, it also indicates that majority of the functional capacity of the human gut virome still remains to be understood.

## Authors and contributions

**KB**: Conception, design, analysis and interpretation of data, original draft preparation, project administration.

**Niharika**, **AG**, **AJ**, **MK**, **MD**, **VS**: Analysis, revisiting the manuscript and final approval and final approval

**SV**: Conception, design, revisiting the manuscript and final approval.

## Acknowledgments

KB acknowledges the support from the DBT, grant no. BT/PR18657/BIC/101/507/2016. SV acknowledges the support from the SERB grant no. JCB/2021/000015.

---

**Figure S1.**
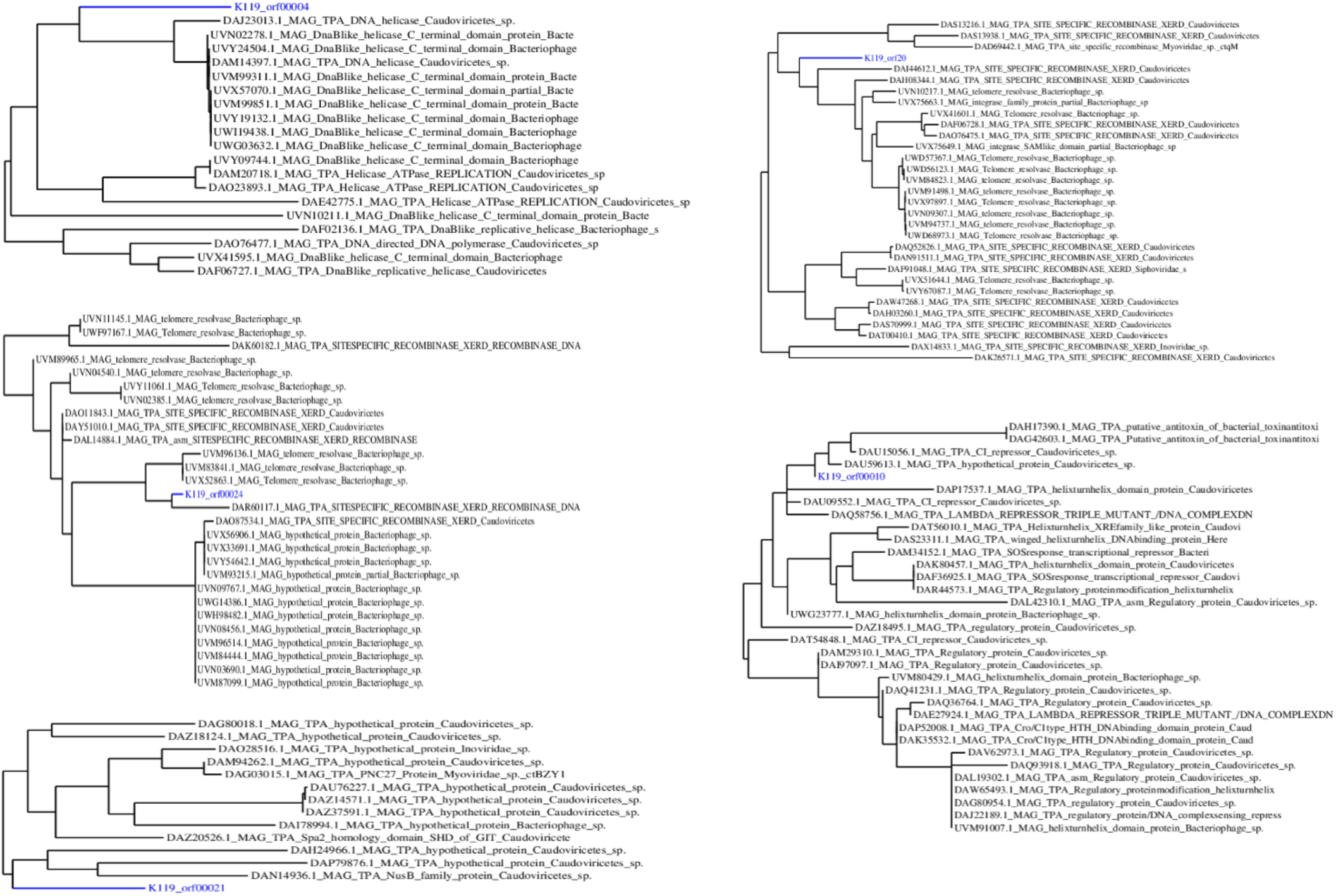
Phylogenetic analysis of the genome, K119_1256_SwiBio_F_0_11, based on five of its predicted ORFs.

**Figure S2.**
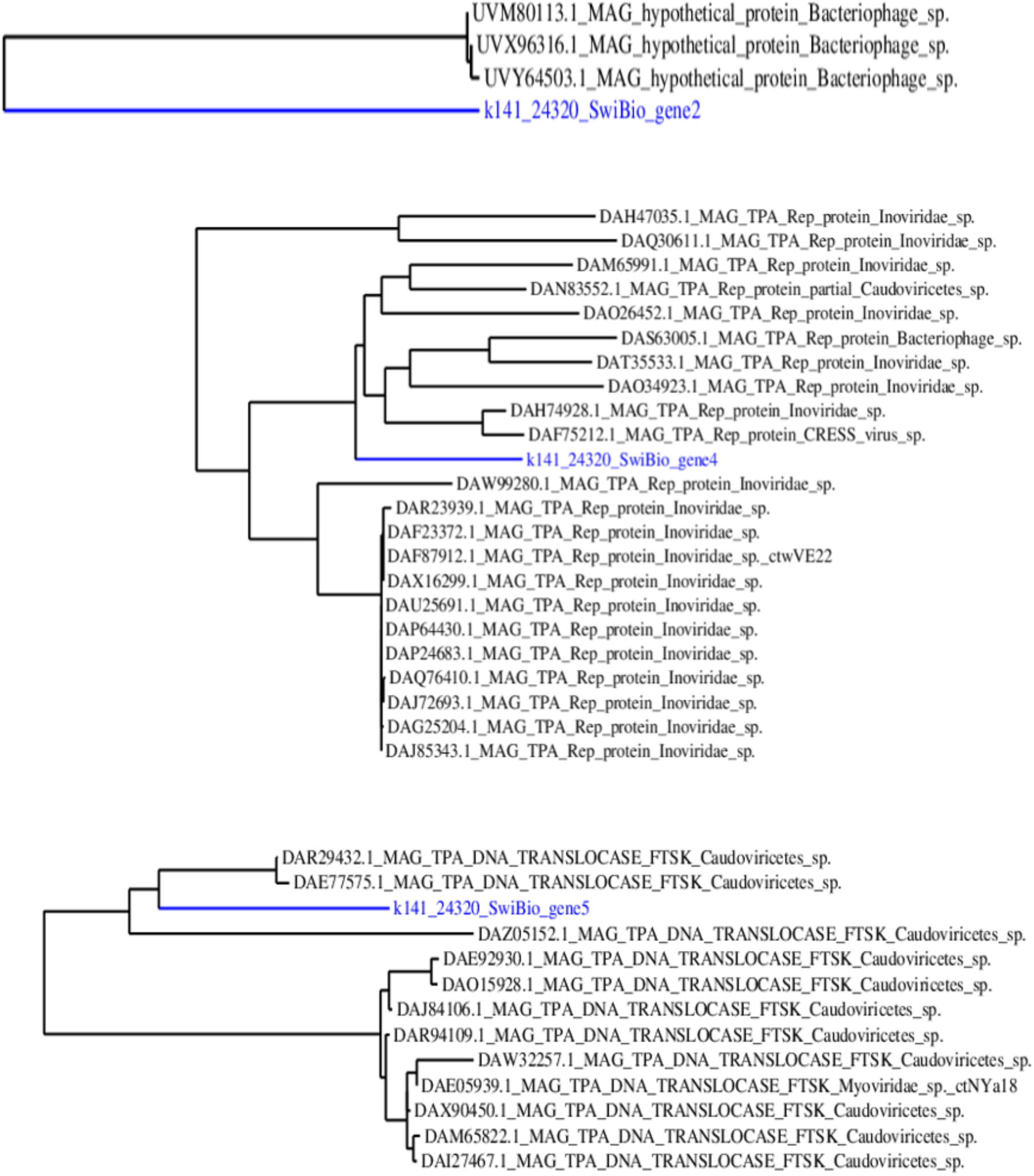
Phylogenetic analysis of the genome, K141_24320_SwiBio_F_0_13, based on three out of five of its predicted genes. Genes 1 and 3 returned no significant hits.

**Figure S3.**
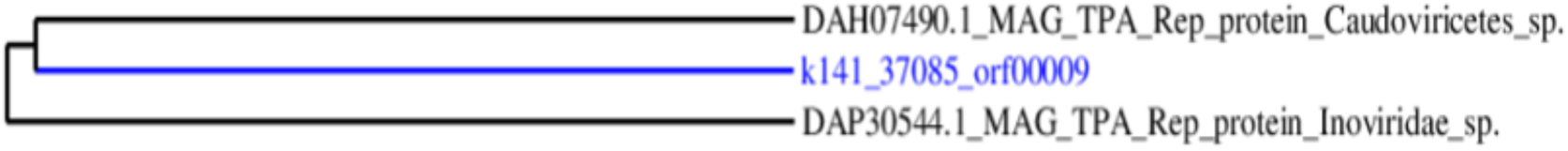
Phylogenetic analysis of the genome, K141_37085_Bac_F_0_13. The other predicted ORFs returned no significant hits.

**Figure S4.**
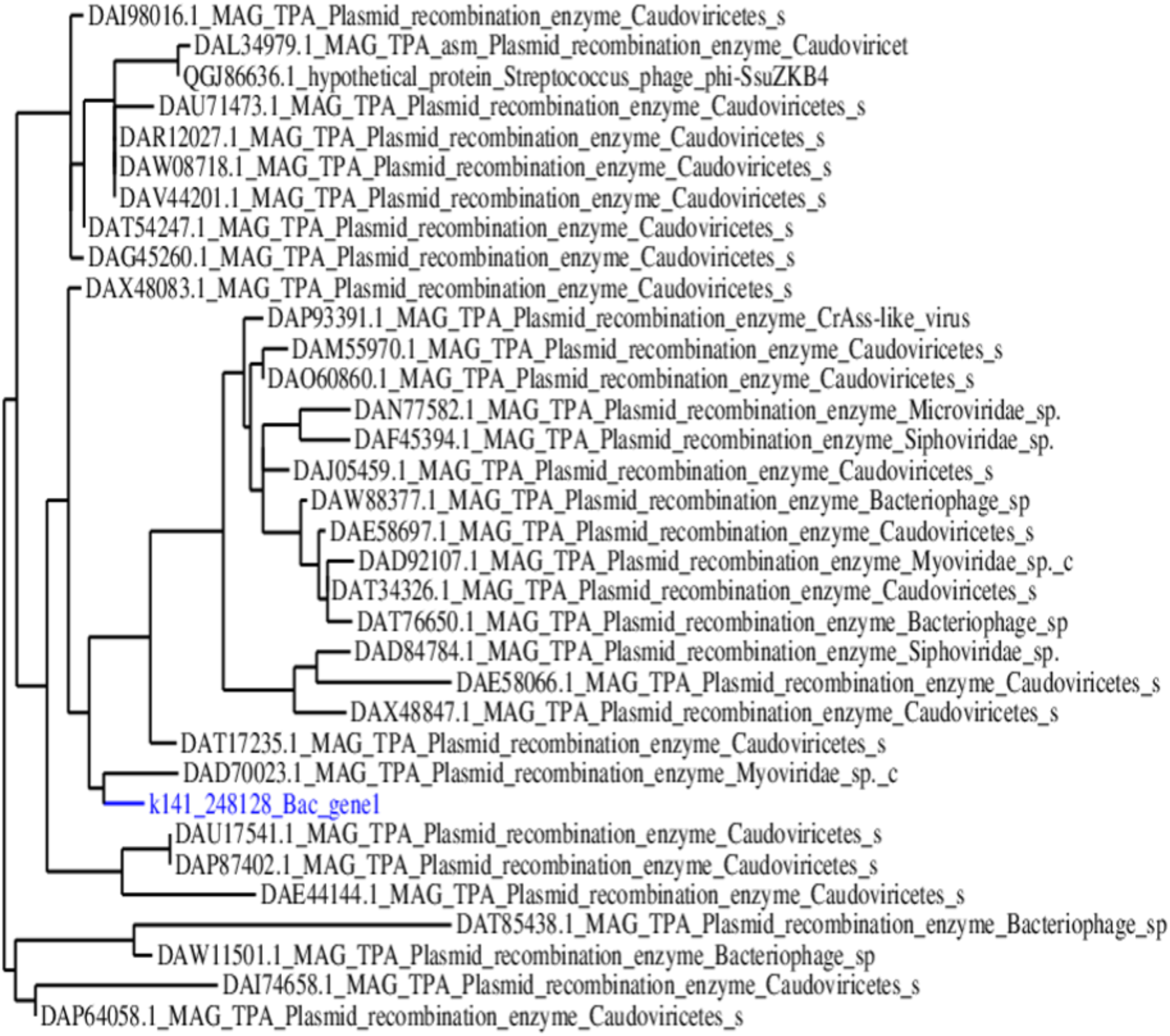
Phylogenetic analysis of the Genome, K141_248128_Bac_F_0_21, based on one of its predicted genes. The other predicted genes returned no significant hits.1

